# Genomic features underlying the evolutionary transitions of *Apibacter* to honey bee gut symbionts

**DOI:** 10.1101/2020.09.30.321786

**Authors:** Wenjun Zhang, Xue Zhang, Qinzhi Su, Min Tang, Hao Zheng, Xin Zhou

**Author notes:** Wenjun Zhang and Xue Zhang contributed equally to this work. Correspondence: Xin Zhou, Department of Entomology, College of Plant Protection, China Agricultural University, Beijing, 100083, China.

## Abstract

The symbiotic bacteria associated with honey bee gut have likely transformed from a free-living or parasitic lifestyle, through a close evolutionary association with the insect host. However, little is known about the genomic mechanism underlying bacterial transition to exclusive adaptation to the bee gut. Here we compared the genomes of bee gut symbionts *Apibacter* with their close relatives living in different lifestyles. We found that despite of general reduction in the *Apibacter* genome, genes involved in amino acid synthesis and monosaccharide detoxification were retained, which were likely beneficial to the host. Interestingly, the microaerobic *Apibacter* species have specifically acquired genes encoding for the nitrate respiration (NAR). The NAR system is also conserved in the cohabiting bee microbe *Snodgrassella*, although with a differed structure. This convergence implies a crucial role of respiratory nitrate reduction for microaerophilic microbiomes to colonize bee gut epithelium. Genes involved in lipid, histidine degradation are substantially lost in *Apibacter*, indicating a transition of the energy source utilization. Particularly, genes involved in the phenylacetate degradation to generate host toxic compounds, as well as other virulence factors were lost, suggesting the loss of pathogenicity. Antibiotic resistance genes were only sporadically distributed among *Apibacter* species, but condensed in their pathogenic relatives, which may be related to the remotely living feature and less exposure to antibiotics of their bee hosts. Collectively, this study advances our understanding of genomic transition underlying specialization in bee gut symbionts.

**Importance:** Investigations aiming to uncover the genetic determinants underlying the transition to a gut symbiotic lifestyle were scarce. The vertical transmitted honey bee gut symbionts of genus *Apibacter* provided an rare opportunity to tackle this, as evolving from family Flavobacteriaceae, they had phylogenetic close relatives living various lifestyles. Here, we documented that *Apibacter* have both preserved and horizontally acquired host beneficial genes including monosaccharides detoxification that may have seeded a mutualistic relationship with the host. In contrast, multiple virulence factors and antibiotic resistance genes have been lost. Importantly, an highly efficient and genomic well organized respiratory nitrate reduction pathway is conserved across all *Apibacter* spp., as well as in majority of *Snodgrassella*, which colonize the same habitat as *Apibacter*, suggesting an crucial role it played in living inside the gut. These findings highlight genomic changes paving ways to the transition to a honey bee gut symbiotic lifestyle.

## Introduction

Bacterial symbiotic association with insect hosts are ubiquitous in nature, conferring traits that enable exploration of new ecological niches (1). Symbiotic bacteria live either intra-or extra-cellularly inside the insect, providing hosts with vital benefits, including nutrients, pathogen resistance and assistance in immunity development (2, 3). Mutualistic symbionts may have various origins, including environmental bacteria or infective parasites (4, 5). Despite distinct evolutionary pathways, the convergent transition to a mutualistic symbiotic lifestyle often involves the loss of virulence factors, degeneration of superfluous functions, and either preservation, or novel acquisition of traits beneficial to the host (6, 7). However, empirical evidence showing the course of transition to a insect symbiotic lifestyle is scarce. Alternatively, comparative genomic analysis of mutualistic symbionts with closely related free-living lineages provides a feasible route to examine the evolutionary phenomena and adaptive mechanisms accompanying the transition to mutualism (8, 9).

Honey bees have simple but specific gut symbiotic bacteria from genera of *Gilliamella, Snodgrassella, Lactobacillus* and *Bifidobacterium* (10). They compose more than 95% of the whole gut community, and the association with these five core bacteria can be dated back prior to the divergence of social corbiculate bees (i.e., honey bee, bumble bee, stingless bee) (11). Although bee gut bacteria are not intracellular, or transovarially transmitted microbes, they are acquired among worker bees in the colony through social contacts (12). Since the establishment of symbiotic association, these bacteria have been subjected to genome reduction and evolutionary adaptation (13). This adaptive transition increases their fitness to the bee gut niche, but also limits the capacity to thrive in other environments (14). As with other symbiotic microbes, honey bee gut bacteria share ancestry with bacterial species carrying varied lifestyles (15). Subsequently, genomic changes are expected to be remarkably different between the transitions from the common ancestry to gut symbionts and to other lifestyles (16). Thus, the honey bee-gut bacteria system provides a promising model to explore the genomic features underlying lifestyle transitions of free-living bacteria to mutualistic gut symbionts. *Apibacter* is currently identified as a genus of honey bee associated bacteria that is prevalent in *Apis cerana, Apis dorsata* and bumble bee species (genus *Bombus*), but only sporadically present in *Apis mellifera* (11, 17). Phylogenetic relationship showed that *Apibacter* spp. formed a monophyletic lineage embedded in the Flavobacteriaceae family, representing the only honey bee gut microbe taxonomy from the phylum Bacteroidetes (17). As a member of the *Chryseobacterium* clade, which is one of the five clades in the Flavobacteriaceae family, *Apibacter* are phylogenetically related to groups of bacteria showing a variety of lifestyles, including environmental free-living (e.g. *Chryseobacterium* genus strains), opportunistic or obligate mammal and bird pathogens (18). To date, only four honey bee associated *Apibacter* strains have been sequenced for whole genomes, allowing the inference of the genomic characteristics, metabolic capacities and cellular features for *Apibacter* (19). However, genetic factors underlying the transition to a bee gut symbiotic lifestyle remain to be uncovered. Additionally, basic biological issues, such as inhabiting location within the honey bee gut, prevalence within natural *A. cerana* populations, and host-specificity remain unknown for *Apibacter*.

In this study, we sequenced 14 *Apibacter* genomes isolated from the Asian honey bee *Apis cerana*. By comparing the genomes of honey bee gut symbiotic *Apibacter* spp. with its close relatives living a variety of lifestyles, we aim to reveal key genomic factors of *Apibacter* to a mutualistic relationship with the honey bee. Moreover, using microscopic visualization, 16S rRNA gene amplicon sequencing and colonization experiment, the spatial habitant, relative abundance in natural *A. cerana* bees, and colonization specificity of *Apibacter* was inferred.

## Results

### The distribution of *Apibacter* in the gut of *A. cerana* and the prevalence among individual bees

The spatial localization of *Apibacter* inside the gut and the distribution through different organs (midgut, ileum, rectum) of the gut remains unknown. Here, we characterized the colonization of *Apibacter* in the gut of *A. cerana* using fluorescence in situ hybridization (FISH) and qPCR. Honey bee symbionts *Snodgrassella* and *Gilliamella* are dominant in the ileum, with *Snodgrassella* colonizing the inner wall of ileum and *Gilliamella* lining on the lumen side of *Snodgrassella* (10). These two bacteria were used as reference coordinates to infer the location of *Apibacter* using species-specific probes (Fig. 1A-D). The spectral images showed that *Apibacter* co-resided with *Snodgrassella* in both ileum and midgut (Fig. 1A, C). The signals were stronger at the inner walls, indicating that *Apibacter* colonized the gut intima as did *Snodgrassella*. *Gilliamella* covered on the top of *Apibacter* and extended into the gut lumen (Fig. 1B, D). *Apibacter* could not be visualized clearly in the rectum, as the rectum was filled with pollen grains with auto-fluorescence under the excitation wavelength. The quantification of the absolute abundances of *Apibacter* in different gut compartments (midgut, ileum, rectum) using qPCR showed that these increased from midgut to rectum, with cell numbers ranging from 1.77 × 10^6^ to 1.87 × 10^7^ (median, Fig. 1E).

**Figure 1.**
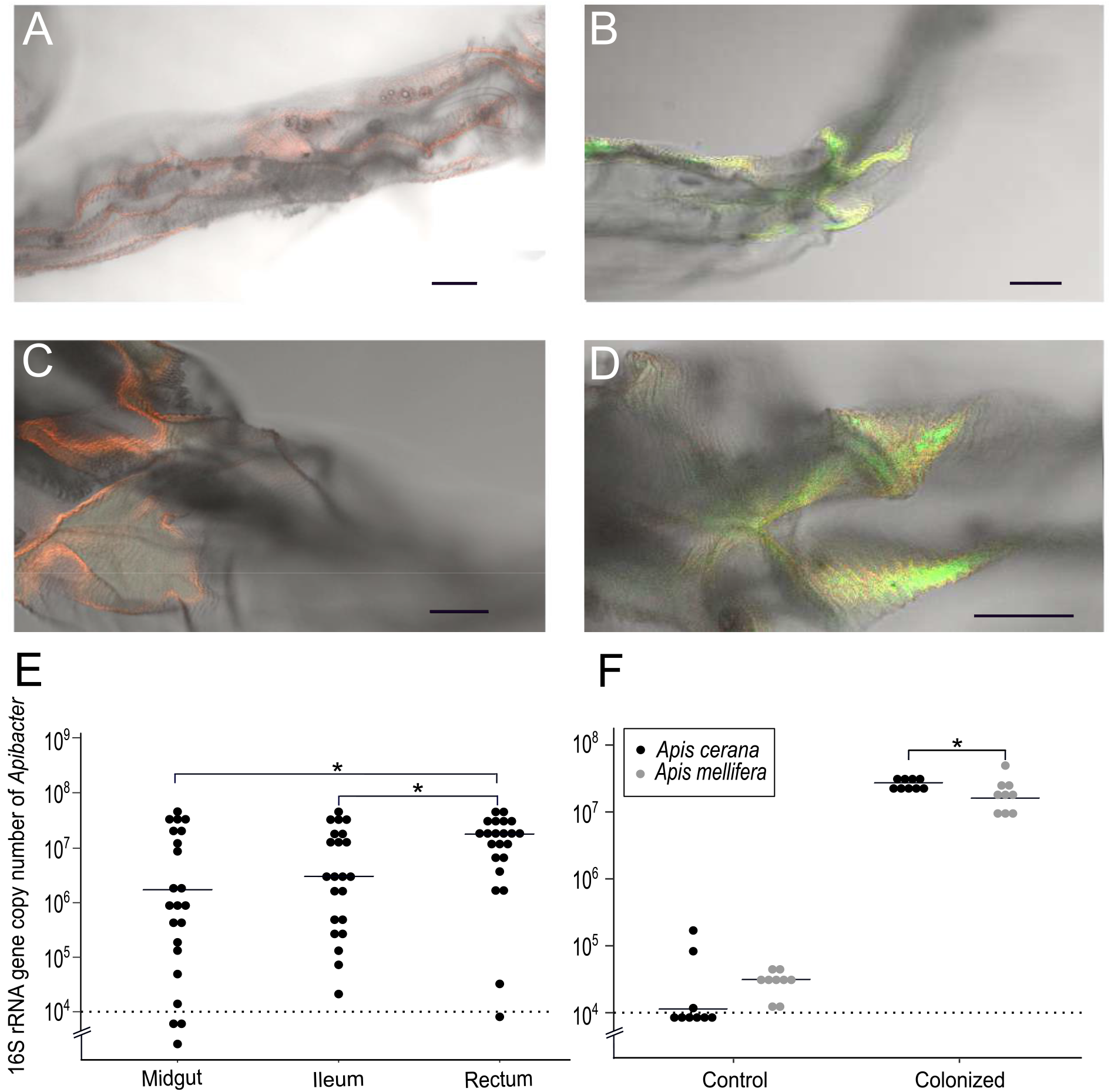
Characterization of the *Apibacter* colonization in honeybees gut. A - D. Localization of *Apibacter* spp. coordinating with *Snodgrassella* and *Gilliamella* within the midgut and ileum of *A. cerana* worker bees. A and B were captures of the ileum. C and D were captures of the midgut. Red fluorescence was the emission of Cy3 fluorophore labeled the *Apibacter* probes and green fluorescence was the emission of Cy5 labeled the *Snodgrassella* and *Gilliamella* probes. The bar represents 100 μm. E. The colonization abundance of *Apibacter* spp. at different intestinal organs in the gut of *A. cerana* worker bees. (n=15 per treatment) F. The abundance of *Apibacter* sp. B3706 isolated from *A. cerana* before and after colonization the whole gut of newly emerged germ free bees of *A. cerana* and *A. mellifera* (n=9). The results of Mann–Whitney U tests (*P < 0.05) are shown.

As *Apibacter* shows sporadically association with *A. mellifera* (11). To test if the *A. mellifera* host immunity exclude the colonization by *Apibacter*, microbe-free bees of both *A. cerana* and *A. mellifera* were inoculated with *Apibacter* sp. strain B3706 isolated from *A. cerana* (see Methods). Six days after inoculation, the cell numbers quantified using qPCR were 2.66 × 10^7^ and 1.64 × 10^7^ (median) in *A. cerana* and *A. mellifera* respectively, both of which were much higher than in the inoculum (< 10^6^), indicating that strain B3706 was able to colonize the guts of both *A. cerana* and *A. mellifera*. However, the cell number in *A. cerana* was slightly, although significantly (Mann-Whitney U test, P < 0.05) higher than that in *A. mellifera*, despite that *A. cerana* had smaller bodysize and gut volume (Fig. 1F), suggesting that strain B3706 was less adapted to the gut of *A. mellifera*. The successfully colonization of *A. mellifera* refuted its immunal rejection to *Apibacter*. Alternatively, interbacterial competition may underlying the low prevalence of *Apibacter* among *A. mellifera* (20).

We further examined whether *Apibacter* and *Snodgrassella* demonstrate interspecific competition in their natural host, which was suspected in a previous study on honey bee gut microbiota (19). The relative abundances of *Apibacter* and *Snodgrassella* were referred based on the gut microbial 16S rRNA gene amplicons sequencing of *A. cerana* worker bees captured from Sichuan, Jilin and Qinghai province in China. The relative abundances of *Apibacter* greatly varied among individual bees and didn’t show obvious inverse relationship with that of *Snodgrassella*, implying no obvious competition between these two co-inhabiting bacteria (Fig. S1).

### Genome characteristics of *Apibacter* and phylogenetic inference

14 strains of *Apibacter* were isolated from worker bees of *A. cerana* from Sichuan, Jilin, and Qinghai provinces in China (Dataset S1). Two genomes (strains B3706 and B2966) sequenced on the PacBio platform were assembled into single circular chromosomes. The remains were sequenced with either Illumina or BGISEQ and were assembled into 12-49 contigs. The genome completeness was evaluated using CheckM (21). All genomes showed a full completeness (Table S2). *Apibacter* strains isolated from *A. cerana* had genome sizes ranging from 2.26 to 2.35 Mb, similar to that from the bumble bees (2.33 Mbp; 22), but smaller than those from *A. dorsata* (2.63–2.76 Mbp; 19) (Table S2). Pathogenic bacteria living a parasitic lifestyle within mammals or birds, e.g. strains from the genera *Riemerella* and *Bergeyella* of Clade C, *Weeksella* and *Ornithobacterium* of Clade E, also have small genome sizes ranging from 2.16–2.44 Mb comparable to other type strains of their clades, suggesting independent genome reductions (Fig. S2 B, Table S2) (23). The low GC content of *Apibacter* was in congruent with the features of animal symbiotic bacteria which have been subjected the mutational bias and weak selections as vertically transmitted through generations with restricted numbers known as transmission bottleneck (Fig. S2 B) (3, 24).

*Apibacter* genomes isolated from *A. cerana* have the pairwise average nucleotide identity values (ANIs) higher than 95.8% (Fig. S3), indicating that they are likely to form a single species. Among the 14 genomes sequenced in this study, six have almost identical ANIs (99.99%) to other isolates (strain B3887, B3883, B3935 comparing to B3889, strains B3918, B3913, B3912 comparing to B3813), such that were excluded from the following analysis to avoid analysis bias due to repeated sampling the same genotype (Fig. S3). Genomic divergence in *Apibacter* isolates is more pronounced between bee species, as the pairwise ANIs are only 85% and 74% in comparisons of *A. cerana* isolates versus those from bumble bees and *A. dorsata*, respectively. Despite the large sequence divergence, genome structure is mostly conserved across the *Apibacter* strains from *A. cerana* and *Bombus*, while a few gene rearrangements and inversions are observed in *Apibacter* genomes from *A. dorsata* (Fig. S4). The conserved genomic structures is in congruent with the observations in *Bartonella apis* from *A. mellifera*, and in *Buchnera aphidicola* from aphids (15, 25).

A maximum-likelihood phylogeny was inferred using a total of 681 orthologous single-copy genes present in all *Apibacter*, related taxa from the family Flavobacteriaceae and one outgroup (*Flavobacterium aquatile* LMG 4008) chosen from family Flavobacteriaceae (Fig. S2 A) (26, 27). Consistent with the previous phylogenetic relationship (17), *Apibacter* species formed a monophyletic group and the isolates from *A. cerana* clustered together (Fig. S2 A). The *Apibacter* clade was sister to the lineage consisting of *Elizabethkingia, Riemerella* and *Chryseobacterium* genera (referred to as Clade C hereafter following Kwong and Moran) (17), most of which are mostly environmental free living bacteria, but also opportunistic pathogenic strains from *Elizabethkingia* and parasitic pathogenic strains from *Riemerella* (18, 28). The *Apibacter* clade together with Clade C formed a sister relationship to Clade E (*Empedobacter, Weeksella*, and *Ornithobacterium*), which mostly consisted of pathogenic strains living environmentally or associated with mammals or birds (29, 30). The fact that *Apibacter* phylogenetically related to bacteria with a complex lifestyles made it a promising model to explore the genomic factors underlying the adapatation to the bee gut environment.

### *Apibacter* undertook a large number of genes loss but preserved specific host beneficial functions

To infer the genomic features responsible for the transition to gut symbionts comparing to other lifestyles, a comparative genomic analysis between *Apibacter* and bacteria of the other two phylogeny clades was conducted. All genes of the bacteria from three clades and one outgroup were clustered into homologous protein families using OrthoMCL (27), and the patterns of the gene gains and losses were inferred by mapping the occurrence of gene families onto the phylogenetic tree using generalized parsimony (31). Based on the gene flux analysis, 601 and 226 protein families were estimated to be lost and gained respectively in the last common ancestor (LCA) of *Apibacter* spp. (Fig. 2A). Protein families of functional categories including ‘Inorganic ion transport and metabolism’, ‘Transcription’, ‘Amino acid transport and metabolism’, ‘Carbohydrate transport and metabolism’, ‘Cell membrane biogenesis’, ‘Lipid transport and metabolism’ (COG category ‘P’, ‘K’, ‘E’, ‘G’, ‘M’ and ‘I’ respectively) were substantially lost in *Apibacter* (9, 9, 8, 8, 7 and 6% of genes with COG annotations respectively (Dataset S2).

**Figure 2.**
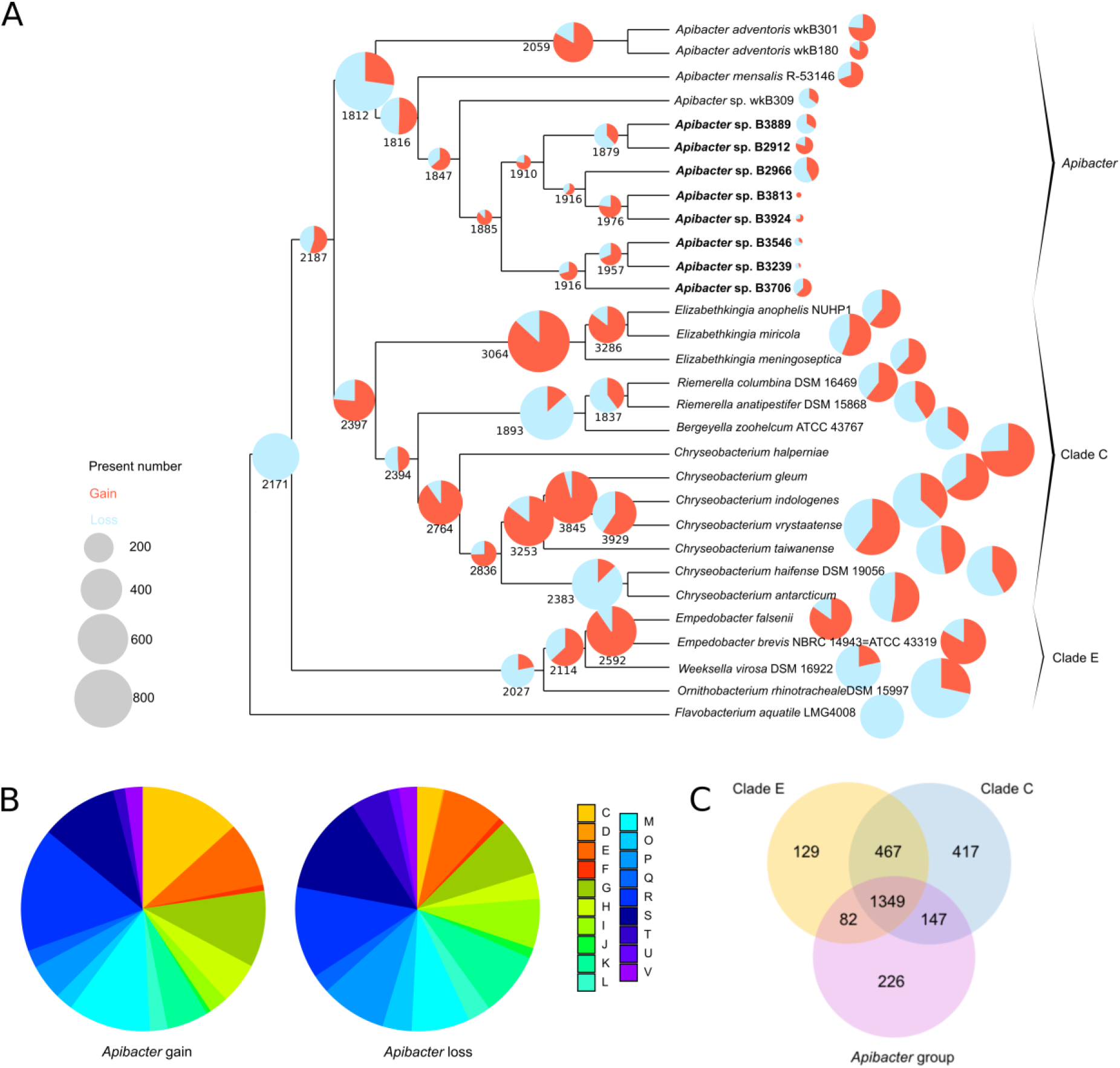
Gene flux analysis and functional classification of niche specifying genes. A. Loss and gain of gene families (red, gain; blue, loss) in gene content since the last common ancestor (numbers in black). The size of the pie chart reflects the amplitude of total gene flux (gain + loss). B. COG functional classification of genes gain and loss in the *Apibacter* genome. See Dataset S2 for complete list of gene families with annotations and COG category abbreviations. C. Venn diagram shows gene family distribution among the three major groups: *Apibacter* group, Clade C and Clade E.

A few bee gut symbionts are capable of degrading the cell walls of pollen granules (*Gilliamella, Bifidobacterium, Lactobacillus*), facilitating polysaccharide utilization for the host (32, 33). Bacteria from the Bacteroidetes phylum confer these functions in human gut (34). However, as a member of Bacteroidetes phylum, *Apibacter* lost most of the genes involved in carbohydrate metabolism. The carbohydrate active enzymes (CAZymes) genes in bacteria from the Bacteroidetes phylum are generally organized into polysaccharide utilization loci (PUL) (35). Compared with members of Clade C and E, *Apibacter* lost most of the PULs. Less than two PULs remained in *Apibacter* that are composed of only tandem *susCD*-like genes, lacking all surrounding CAZyme genes (Dataset S3). The PUL residues in the genome of *Apibacter* isolates from *A. cerana* and *Bombus* are conserved in protein sequences (>87% similarity). One PULs in the genomes of *Apibacter adventoris* wkB301 from *A. dorsata* showed similarity to the PULs from *A. cerana* and bumble bee (>70% similarity), while the other three are unique to isolates from *A. dorsata* (Dataset S3).

The gained protein families were mostly associated with functional categories of ‘Energy production and conversion’, ‘Cell membrane biogenesis’, ‘Carbohydrate transport and metabolism’ and ‘Amino acid transport and metabolism’ (COG category ‘C’, ‘M’, ‘G’ and ‘E’, 13, 11, 10 and 9 % of genes with COG annotations respectively, Dataset S2). Exemplified by bee associated *Snodgrassella*, symbiotic gut bacteria are capable of synthesizing amino acids for hosts and other co-occurring symbionts (7, 13). Consistently, *Apibacter* have preserved all genes underlying amino acid synthesis, which are inherited from the LCA. On the contrary, the parasitic pathogens from genera *Bergeyella*, *Weeksella*, *Riemerella* and *Ornithobacterium*, which have also been subjected to independent genome reductions, lost substantial proportions of their ancestral amino acid synthesis genes (Dataset S4). Moreover, genes beneficial to the host were particularly preserved. For instance, the mannose-6-phosphate isomerase encoded by *manA* is responsible for degradation of the toxic mannose for the host (36). This gene is lost in all strains of Clade C and E, but is preserved by all *Apibacter* strains, indicating that mannose detoxification is an important mutualistic trait in *Apibacter* (Dataset S5). *Apibacter* also particularly gained genes involved in L-arabinose and xylose degradation (*araB, araD* and *xylB*, Dataset S2). A BLASTP search against the non redundant protein sequences (nr) database showed that the AraB and AraD of *Apibacter* had best hits to distantly related bacteria from other classes within Bacteroidetes, while the XylB showed a best hit to bacteria from the phylum Actinobacteria, suggesting a horizontal acquisitionof these genes. These two monosaccharides are main constituent components of hemicellulose and pectin and are also toxic to the bee host (37). The acquisition of genes for the degradation of the toxic monosaccharides potentiates *Apibacter* with the ability to utilize the pollen hydrolysis products, at the same time enabling monosaccharide detoxification for the host.

### Respiratory nitrate reduction pathway was conserved in *Apibacter* group

In order to identify genomic characteristics related to the specifically transition to the bee gut symbionts, the protein family contents in the last common ancestors (LCA) of three clades were inferred and compared. Differentiated from a pangenomes analysis that involves protein families only sporadically presented in few isolates due to the strain level diversity (38), the protein families in the common ancestor are mostly conserved in the majority members of the clades. Results showed that 1,349 core genes were shared by the LCA of all three clades (Fig. 2C). 226 protein families were specific to the LCA of *Apibacter* clade, overrepresented by the gene functionalities of nitrate assimilation, glycerol catabolism, molybdate ion transmembrane transporter activity, L-arabinose catabolism, cysteine-type peptidase activity (Hypergeometric test, p < 0.05, Dataset S6). A closer inspection showed that the genes involved in the respiratory nitrate reduction (NAR) pathway co-located in the genomes and were conserved among all *Apibacter* isolates (Fig. 3, Dataset S5). However, one of the NAR pathway genes, *narI*, encoding the membrane biheme b quinol-oxidizing γ subunit of the nitrate reductase was missing. Functioning in its place was a protein shared a highly conserved domain with Rieske protein (cl00938 in NCBI Conserved Protein Domain Family) that located inside the *Apibacter* NAR gene cluster. This Rieske protein is an iron-sulfur protein (2Fe-2S), with a function in the transferring of electrons from the quinone pool, same as that of NarI (39, 40). The missing of *narI* in the nitrate reductase complex was previously reported in halophilic archaea, which represents an ancient respiratory nitrate reductase (41, 42).

**Figure 3.**
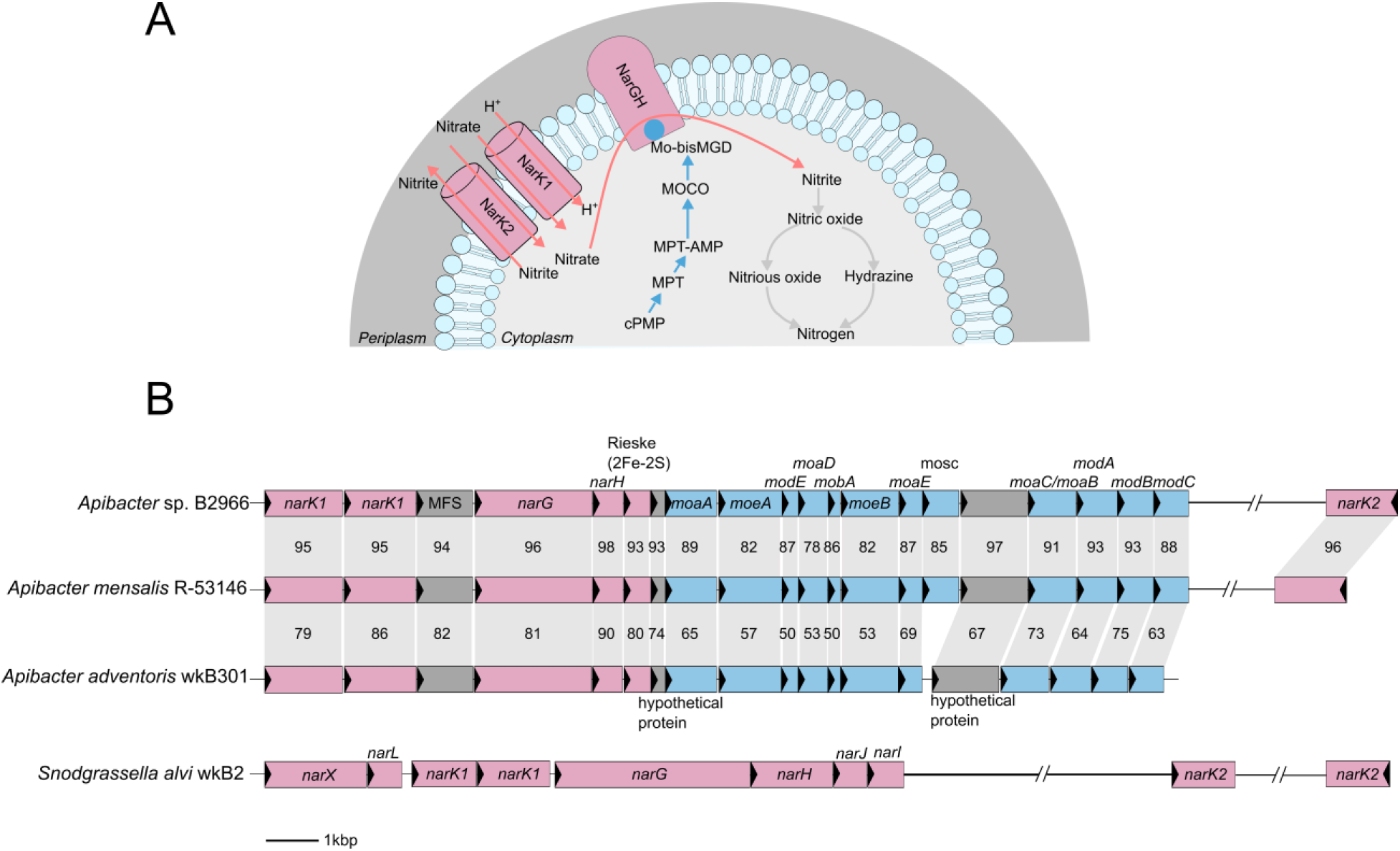
The respiratory nitrate reduction pathway is conserved in the *Apibacter* group, and is also possessed by *Snodgrassella*. A. Schematic diagram shows that this together with molybdate cofactor synthesis are conserved accoss the *Apibacter* group (pink and blue arrows), and nitrite detoxification (grey arrows) is absent in the *Apibacter* group but possessed by genomes in Clade C/E (genes distribution are provided in Dataset S5). B. Genomic regions of type strains of *Apibacter* from *A. cerana*, bumble bee and *A. dorsata* encode genes involved in respiratory nitrate reduction. Vertical grey blocks connect homologous genes among type strains, with numbers representing the percentage of sequence similarities. Genes in pink encode nitrate reductase and transporters. Genes in blue encode molybdate cofactor synthesis. Genes in grey are hypothetical genes or genes that are not directly related.

Molybdenum cofactors (Moco) are required for the activity of NAR nitrate reductase (43, 44). The genes involved in the biosynthesis of the Moco were also enriched and are conserved across *Apibacter* genomes, located adjacent to the NAR related genes, although one Moco synthesis gene (*mosC*) and one nitrate transporter was missing in the two *A. dorsata* isolates (Fig. 3). The NAR related genes were highly conserved in amino acid sequences among genomes isolated from the same bee hosts (>94% similarity). Isolates from *A. cerana* are more similar to the one from bumble bee than those from *A. dorsata* (Fig. 3). Another defining characteristic of this NAR pathway is the presence of three copies of the *narK* nitrate transporter genes in *Apibacter* (Fig. 3B) (45), implying high efficiency of nitrate respiration in *Apibacter*. A phylogeny based on the NarG and NarH protein showed that *Apibacter* spp. symbionts are related to bacteria of *Flavobacterium* genus and other members in the Bacteroidetes phylum (Fig. S5). Most species of Flavobacteriaceae contain single copy of these genes. This phylogenic inference implied that these genes were most likely inherited from the ancestry.

In contrast, the NAR pathway was absent in bacteria from Clade C and E, which was congruent with their aerobic nature. It is worth noting that parasitic pathogens from *Riemerella* and *Ornithobacterium* also lack the NAR operon (28, 30), despite the fact that they are microaerophilic, like *Apibacter* (Fig. 3, Dataset S5). These results imply that the NAR pathway might be particularly beneficial to gut commensal bacteria, which prompted us to survey the presence of NAR pathway in other honey bee gut bacteria. Interestingly, the NAR pathway was conserved in the majority of *Snodgrassella* strains although showed a varied structure from that in *Apibacter*. The *narGHJI* genes encoding the four subunits of nitrate reductase were intact in *Snodgrassella* with a similar pattern to those in *E. coli* (Dataset S5) (43), while genes involved in Moco synthesis located separately from the nitrate reductase genes (Fig. 3B). In contrast, the NAR genes were mostly absent in the other four core bee gut bacterial phylotypes (*Gilliamella, Bifidobacterium, Lactobacillus* Firm4 and Firm5) (Dataset S5), leading to the hypothesis that the NAR pathway might be generally required by microaerobic bacteria inhabiting the gut epithelium.

### *Apibacter* lost ancestral gene families related to pathogenicity and antibiotic resistance

The LCA of Clade C and Clade E had 467 protein families in common, representing unique functionalities to a variety of lifestyles, but not involved in bee gut symbiosis. These protein families overrepresented in functionalities of fatty acid beta-oxidation, nickel cation binding, phenylacetate catabolic process, potassium-transporting ATPase activity, histidine catabolic process, beta-galactosidase activity and urease activity (Hypergeometric test, p < 0.05, Dataset S6). Gene families associated with these functionalities were examined across all analyzed genomes by performing a TIGRFAM Hidden Markov Model (HMM) search (46) or BLASTP against the Uniprot Database (47). Genes in the histidine degradation to glutamate and formamide (Hut pathway) were completely lost in all *Apibacter* isolates (Fig. 4, Dataset S5). Histidine biosynthesis is one of the most energy consuming processes in bacteria, such that the degradation of histidine as carbon and nitrogen sources is strictly regulated (48). Oxygen is required for the activation of the Hut operon (49). Given that the bee gut is mostly anoxic, the Hut pathway is highly likely to be malfunctioning in *Apibacter* and is susceptive to be lost. In contrast, histidine degradation is important for pathogens to recognize eukaryotic hosts and to activate virulence factors (50). The Clade C and E included important pathogenic bacteria to mammals and birds, among which histidine degradation may important features for host infection. The acyl-CoA dehydrogenase (FadE) catalyzing the first step in the lipid beta-oxidation cycle was absent in all *Apibacter* isolates, suggesting that fatty acid is not a feasible energy source. In addition, fatty acid oxidation generates organic electron donors (reduced flavin adenin dinucleotide) via the acyl-CoA dehydrogenase for nitrate respiration(51). The loss of *fadE* indicates that fatty acid can not serve electrons to the nitrate respiration in *Apibacter* strains. Ring 1,2-epoxyphenylacetyl-CoA and 2-oxepin-2(3H)-ylideneacetyl-CoA are two intermediate substrates generated in the phenylacetate degradation that are toxic to animal host (52, 53). Genes (*paaACD/paaG*) encoding for enzymes responsible for the catabolic transversion to these two toxic substrates are absent in *Apibacter*, suggesting the loss of pathogenicity along the transition to the gut symbiosis (Fig. 4). Urease is also an important virulence factor for many pathogenic bacteria (54). As a metalloenzyme, urease requires nickel for its catabolic activity (55). Both urease and nickel cation binding protein families were overrepresented in the Clade C and E shared ancestral genomes, but lost by *Apibacter* (Dataset S6).

**Figure 4.**
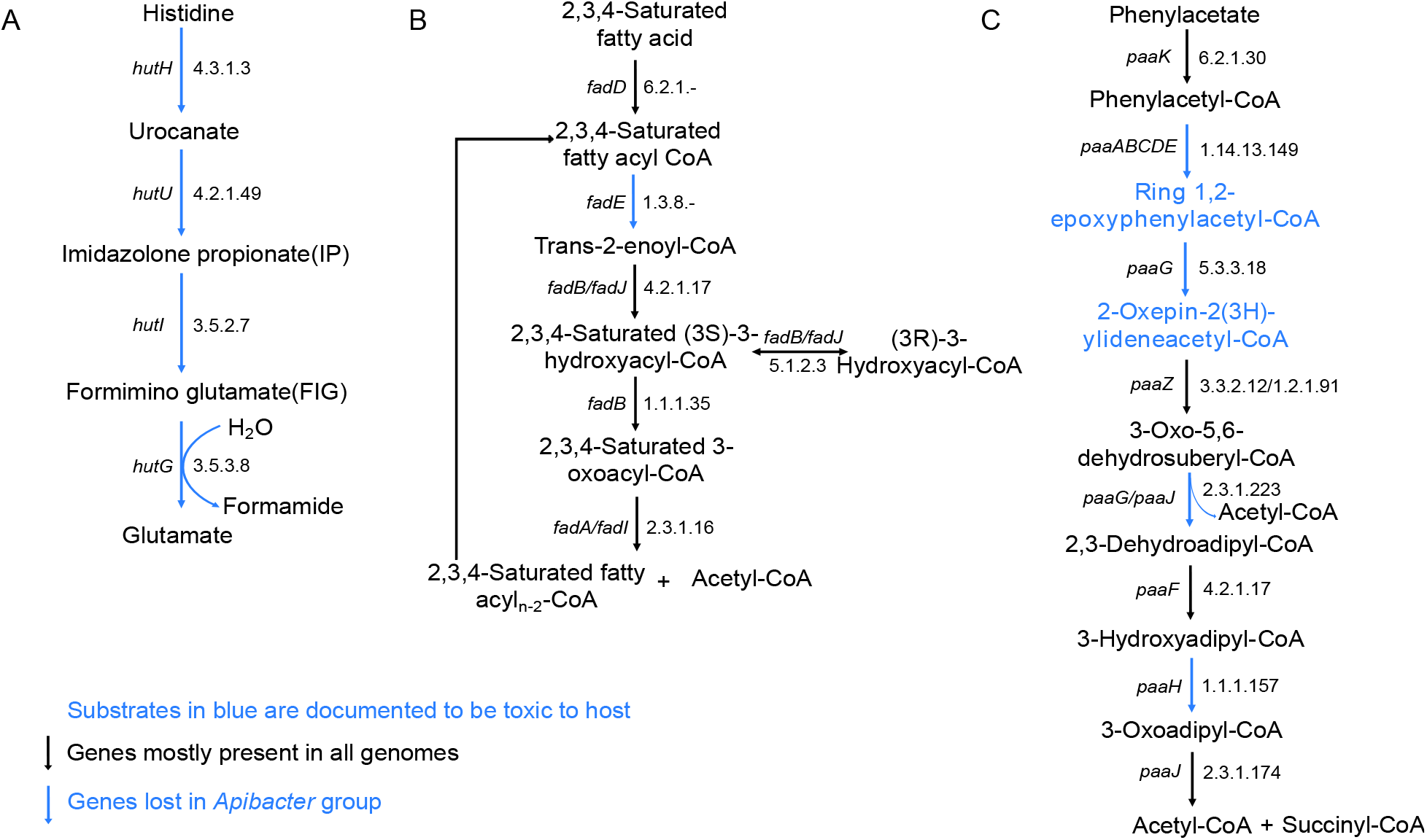
Metabolic pathways that are present in genomes of the Clade C and Clade E, but incomplete among genomes of *Apibacter* group. A. Histidine degradation pathway; B. Fatty acid beta-oxidation pathway; C. Phenylacetate oxidation pathway. Black arrow, genes mostly present in genomes from three groups; Blue arrow, genes are absent among all genomes in *Apibacter* group, but possessed by genomes in Clade C and E (gene distributions are provided in Dataset S5). Substrates in blue in phenylacetate degradation are documented to be toxic to host.

To identify other pathogenicity related gene families that are prevalence in the relative bacteria but lost by *Apibacter* spp., which may represent key virulence factors to honey bee, genomes of the three clades were queried against the virulence factor database (VFDB) (56). A total of 37 virulence factors involved in host cell adhesion and invasion, secretion systems and effectors, toxin production and iron acquisition were identified. Interestingly, the chaperon/usher (CU) pathway and the extracellular nucleation-precipitation (ENP) pathway involved in the host cell adhesion and invasion, and one DNase I genotoxin were absent in almost all *Apibacter* genomes (with the exception of one CU pathway identified in *Apibacter* sp. B2912), but were dominantly distributed among genomes from Clade C and E (Dataset S7). The CU pathways responsible for pili formation are known to be virulence factors for Gram-negative pathogens (57). Particularly, gene encoding products in the CU pathway were documented to be crucial for the urinary tract infection (58, 59), which is in line with the fact that multiple bacteria in Clade C and E (e.g. bacteria from genus *Empedobacter* and *Elizabethkingia*) are urinary infective pathogens (60, 61). The ENP pathway is responsible for the curli production related to host adhesion and inflammatory inducing (62). The DNase I was identified as a polyketide synthetase (PKS) that showed high protein sequence similarity to that from bacteria of family Enterobacteriaceae. PKS genes located in the genomic island and are susceptible to transmission (63). Interestingly, the PKS has also been identified to involved in the symbionts mediated protection against eukaryotic cells including fungus and parasitoid predators in insects (64–66). These virulence factors represent key pathogenicities for pathogenic bacteria, and may confer potential deleterious effects to the bee hosts.

Antibiotic resistances are promiscuous for bacteria in the Flavobacteriaceae family, causing difficulties in the treatment of their infection (18). Referring to the CARD database, 13 antibiotic resistance genes that confer resistance to beta-lactam, fluoroquinolones, tetracyclines and glycopeptides were identified in genomes of Clade C and E (Fig S6). These resistance genes were absent in the *Apibacter* group, except that a lincosamid resistance gene identified in *A. adventoris* wkB301. These results may be explained by the fact that *A. cerana* bee gut microbes are less exposed to antibiotics.

## Discussion

Combining FISH and colonization experiments, we revealed the colonization specificity of *Apibacter* and its distribution in the bee gut. Comparative genomic analyses of 30 genomes from the *Flavobacteriaceae* family, including 8 newly sequenced *Apibacter* genomes from this study and publicly available genomes for the outgroups, helped us to characterize gene signatures underlying the lifestyle transition and adaptation to a lifestyle as bee gut symbionts.

FISH visualization indicates that *Apibacter* cohabit with *Snodgrassella* and colonize the epithelium of the bee gut. As core members of gut bacteria in *A. cerana*, both *Apibacter* and *Snodgrassella* are microaerophilic, sharing nutritional sources (67). We showed that *Apibacter* isolated from *A. cerana* were able to colonize *A. mellifera* suggesting that host incompatibility is probably not the constraining factor responsible for the rarity of *Apibacter* in *A. mellifera*. However, it is not yet possible to examine inter-host competition between *Apibacter* isolated from different honey bee species, because isolates from *A. mellifera* were not available to us.

Comparative genomic analysis revealed gene functions potentially associated with the adaptation to bee gut niche. In a typical symbiotic system, host-level benefits are considered crucial for the establishment of a mutualistic relationship (4, 6). *Apibacter* retained the mannose catabolic gene, which was responsible for monosaccharide detoxification in the honey bee therefore broadening food choice for the host (36). Moreover, genes involved in amino acid biosynthesis are preserved in *Apibacter* spp., at a background of overall genome reduction, which echoes those previously reported in other bee gut symbionts (13).

Polysaccharide utilization is a prominent property encoded by bee gut symbionts including *Gilliamella, Bifidobacterium* and *Lactobacillus* (32, 68, 69). However, relevant genes are substantially lost in the *Apibacter* group. Interestingly, the core bacterial species *Snodgrassella* that cohabit with *Apibacter* along the *A. cerana* gut epithelium also lack the capacity to utilize polysaccharides (13). We speculate that polysaccharides might be limited in the niche that they share.

The gut lumen is mainly anaerobic (67). However, oxygen can diffuse from the intestinal epithelium cells and create a microaerobic environment for facultative anaerobes (67, 70). A previous study found both cytochrome *bd* and *cbb*_3_ in the *Apibacter* genome, which were presumably involved in microaerobic respiration (19). In the present work, we identified an anaerobic respiration NAR pathway that was conserved within the *Apibacter* group and in the cohabiting *Snodgrassella*, but absent from the other four core bee gut bacteria species. These observations suggest that the NAR pathway might be important for the microbiome to colonize bee intestinal epithelium. Such respiratory flexibility might enable *Apibacter* to survive altered oxygen tensions. This finding is congruent with the observation in mouse *E. coli*, where they require both microaerobic and anaerobic respirations for successful colonization (71). A further study proved that the NAR pathway played a key role in *E. coli* colonization of the mouse gut, because the NarG mutant showed colonization deficiency for both commensal bacteria and pathogenic *E. coli* (72). These results are in line with the observation that nitrate reduction could facilitate the growth of gut microaerobic bacteria at low oxygen conditions (73). Therefore, we conclude that the NAR operon is an important genetic signature for *Apibacter* adaptation to the bee gut.

In conclusion, combining molecular and colonization experiments, for the first time, we visualized and quantified the distribution of *Apibacter* spp. inside the bee gut, and proved that *Apibacter* isolates of *A. cerana* could survive in *A. mellifera*. Genomic comparisons with relatives living on other lifestyles revealed that host beneficial traits and respiratory nitrate reduction (NAR pathway) were key functionalities for adaptation to the bee gut environment.

## Materials and Methods

### Sample collection and *Apibacter* isolation

The worker bees of *Apis cerana* were collected from Sichuan, Jilin and Qinghai Provinces in China (Dataset S1). 3 individuals from Sichuan, 1 individual from Jilin and 2 individuals from Qinghai were dissected. The guts were homogenized and frozen in glycerol (25%, vol/vol) at – 80°C. Frozen stocks of homogenized guts were streaked out on heart infusion agar (Oxoid) or Columbia agar (Oxoid) supplemented with 5% sheep blood (Solarbio). The plates were incubated at 35°C in 5 % CO_2_ for 2–3 days. Colonies were screened by PCR using universal primers 27F and 1492R for the16S rRNA gene and Sanger sequencing.

### Fluorescence *in situ* hybridization (FISH) microscopy

Ten adult worker bees of *A. cerana* were collected from a single colony in Beijing, China in May 2018. The whole guts of bees were dissected and frozen in RNAlater (Qiagen) at −80 °C. The FISH protocol was adapted from Yuval et al. (74). In brief, the guts preserved in RNAlater were fixed in the fixative solution (Ethanol: Chloroform: Acetic acid = 6:3:1) and kept overnight at room temperature, followed by rinsing with 1 × PBS. The guts were then treated with 1 mg/mL proteinase K solution at 56 °C for 20 minutes, followed by rinsing with 1 × PBS. The guts were incubated with 6% H_2_O_2_ ethanol solution for 2 hours, followed by rinsing twice with 1 × PBS. Finally, the guts were permeated with 0.1% Triton for 2 hours, followed by rinsing for 2 or 3 times in 1 × PBS buffer.

The genus specific FISH probe targeting the *Apibacter* 16S rRNA genes was used as described previously by Weller et al. (75) (Table S1). The specific probes for the genera of *Snodgrassella* and *Gilliamella* were adapted from those used by Martinson et al. (10) (Table S1). The *Apibacter* specific probes were labeled with Cy3 fluorophore, which the probes for *Snodgrassella* and *Gilliamella* were labeled with Cy5 fluorophore (Table S1). Probe hybridization was performed overnight at 37 °C, followed by rinsing in 1 × PBS for 3 times. Spectral imaging was used to visualize gut sections on a ZEISS LSM780 confocal microscope because of its ability to produce optical sections with reasonable accuracy. And at the excitation wavelength and emission wavelength of 561nm and 633nm, the Cy3 and Cy5 probe-labeled strains showed red and green fluorescence, respectively. Autofluorescence was assayed for each tissue type as the negative control.

### Estimation of bacterial abundance using qPCR

15 worker bees were collected from 2 colonies at the same apiary in Beijing, China in September 2018. Samples were stored at −80°C. Gut segments (i.e., midgut, ileum, and rectum) were dissected and separated. DNA was extracted using the CTAB method following Powell et al. (12). The 27F and 355R universal 16S rRNA primers were employed for all bacteria and *Apibacter*-specific primers Apiq9-F and Apiq9-R were designed in this study (Table S1). The qPCR was performed on a LightCycler 480 (Roche Applied Science, Indianapolis, IN) using the Roche SYBR Green I Master mix. The qPCR reaction was set up as the following: 95°C for 5 min and 40 cycles of three-step PCR at 95°C for 10s, 55°C for 30s and 30s at 72°C with a melting curve observed at the end of each run. Standard curve was determined using a 1:5 series dilution of the *Apibacter* genomic DNA extract. 3 technical replicates were employed for each sample. Significance in differences between samples was determined using Mann-Whitney (Wilcoxon-rank) nonparametric U tests.

### *In vivo* colonization experiments

Microbiota-depleted bees were obtained following Zheng et al. (67). Late stage pupae of both *A. cerana* or *A. mellifera* were removed from brood frames and incubated for 24-36 h in sterile plastic bins at 35 °C and 75% humidity. Newly emerged bees were kept in cup cages provided with sterilized sucrose syrup (0.5 M). For inoculation, a bacteria strain of the *Apibacter* sp. (B3706) was grown on the heart infusion agar (Oxoid), which was supplemented with 5% sheep blood from glycerol stocks, for two days. Cultivated strains were scraped and suspended in 1 × PBS to reach an OD_600_ of 1.0. Batches of 25 bees were placed in a 50 ml conical tube, and 50 μl sucrose syrup was added. The tube was rotated gently so that the syrup was coated on the surface of bees. The tube was rotated again after the addition of 50 μl of the *Apibacter* sp. B3706 suspension (~2 × 10^6^ cells per bee). The bacteria were inoculated into the bee guts as a result of auto- and allogrooming. Inoculated bees were reared in cup cages. Three replicate enclosures were set up for both *A. cerana* or *A. mellifera* with 20 bees in each cage. The inoculated bees were fed with sterilized sucrose syrup throughout the experiment. Colonization levels were determined at day 6 using qPCR as described previously.

### Genome sequencing, assembly, annotation and 16S rRNA gene sequencing

Genomic DNA of *Apibacter* isolates was extracted using a bead-beating method as previously described (12). Two strains (*Apibacter* sp. B2966 and B3706) were sequenced with the PacBio RS (PacBio) at Nextomics Biosciences Co. Ltd., China, and assembled using the OLC algorithm of the Celera (76). Three strains (*Apibacter* sp. B3239, B3546 and B2912) were sequenced with a BGISEQ-500RS (BGI) at BGI-Qingdao, China. The other nine strains (*Apibacter* sp. B3813, B3883, B3887, B3889, B3912, B3913, B3918, B3924 and B3935) were sequenced with a HiSeq X-Ten (Illumina) at Novogene Co., Ltd, China. All strains except for *Apibacter* sp. B2966 and B3706 were assembled with *SOAPdenovo-Trans* (version 1.02, −K 81 - d 5 -t 1 -e 5 for 150PE reads; -K 61 -d 5 -t 1 -e 5 for 100PE reads) (77), *SOAPdenovo* (version 2.04, only for 150PE reads) (78) and *SPAdes* (version 3.13.0, -k 33,55,77,85) (79). The quality trimmed reads were mapped back to the assembled contigs using minimap2-2.9 (80) to examine assembly quality. The bam file generated by *samtools* (version 1.8) (81) and the assemblies were processed by *BamDeal* (https://github.com/BGI-shenzhen/BamDeal, version 0.19) to calculate and visualize sequencing coverage and GC contents of assembled contigs. Spurious contaminants, contigs with low depths of coverage or abnormal GC contents were removed from the draft genome. Assembled genomes were then annotated using PROKKA 1.13.3 (82).

28 bees were captured from 3 provinces in China and the gut microbiome composition was examined using 16S rRNA gene amplicon sequencing (16 of Sichuan, 6 of Jilin, 6 of Qinghai). The gut was dissected and the genomic DNA was extracted using a bead-beating method as previously described (12). Universal primers 340F (5’-CCTACGGGAGGCAGCAG-3’) and 533R (5’-TTACCGCGGCTGCTGGCAC-3’) were used to amplify the V3 region of 16S rRNA gene and sequencing was performed on the BGISEQ-500 2×150 bp platform at BGI-Qingdao, China. Obtained reads were demultiplexed referring to the barcode sequences, trimmed to 100bp. Taxonomy classification was assigned by 97% similarity cutoff using the MOTHUR v.1.40.5 pipeline5 (83).

### Comparisons of genome structure, genome divergence, and gene contents

The contigs of each assembly were re-ordered according to the single circular genome of strain B3706 using the ‘Contig Mover’ tool of Mauve version 2.4.0 (84, 85). Pairwise average nucleotide identity (ANI) was calculated with JSpeciesWS (86) using the BLASTN algorithm (ANIb). Genome completeness was estimated with CheckM (21), which was available at KBase online (87) using recommended parameters. The genome structures were compared using the R-package genoPlotR (88).

### Phylogenetic inference

Gene orthology was determined using OrthoMCL (27) for all genomes used in this study. All steps of the OrthoMCL pipeline were executed as recommended in the manual and the mcl program was conducted using parameters ‘-abc -I 1.5’.

Protein sequences of the identified single-copy orthologous were aligned using Mafft-linsi (89). Alignment columns only containing gaps were removed and the alignments were concatenated. The phylogeny was inferred using RAxML v8.2.10 (26) with the PROTGAMMAIJTT model and 100 bootstrap replicates.

### Gene flux analysis

Gene gain and loss analyses and inferences of gene contents of LCAs (last common ancestors) were conducted using Count (31). Standard methods used in previous works were employed in the present study (15). For each gene family, Wagner parsimony with a gene gain/loss penalty of 2 (90) was used to infer the most parsimonious ancestral states. Parameter choices followed a previous publication (91).

### Analysis of functional gene contents

Gene contents were categorized based on COG and eggNOG (92) functions. Clade specific protein families were assigned with gene ontology (GO) terms and enrichment analysis was performed using OrthoVenn (93). For gene families of interest, BLASTP(94) was used to query against the NCBI’s nr database and UniProt (47). TIGRFAM Hidden Markov Model (HMM) (46) was also used to search for protein families of specific functions. The phylogenies of NarG and NarH were produced from protein sequences obtained by BLASTP against NCBI’s nr database with default parameters. Hits were aligned with *Apibacter* sequences using Mafft-linsi. Alignments were then trimmed with trimAL (95) for sites with over 50% gaps and phylogenetic trees were inferred using RAxML PROTGAMMALG with 100 bootstraps as described previously (96).Carbohydrate-active enzyme (CAZymes) gene families were identified for all analyzed genomes using the command-line version of dbCAN (Database for automated

Carbohydrate-active enzyme Annotation) (97), following authors’ instruction. The PULs (Polysaccharide utilization loci) were identified using the TIGRfam (98) and Pfam (99) models, following Terrapon et al. (100). Annotation of virulence factors and classification the virulence factors were based on VFDB (Virulence Factors of Bacterial Pathogens)(56). All genomes were blast against the classified virulence factors from VFDB. Antibiotic resistance genes were identified by querying all the genomes against the Comprehensive Antibiotic Resistance Database (CARD) (101).

## Acknowledgements

The work was funded by the Program of Ministry of Science and Technology of China (2018FY100403) and National Natural Science Foundation of China (31772493).

## Data Accessibility

The *Apibacter* genome sequences in this study are available under NCBI Bioproject accession PRJNA578212.

The 16S rRNA used in this study are available under NCBI Bioproject accession PRJNA663994.

## Supplementary Datasets

Dataset S1. List of genome sequence deposition and strain collection sites.

Dataset S2. List of *Apibacter-specific* gene families gain and loss and COG category abbreviations.

Dataset S3. Distribution of CAZy genes according to three groups and PULs similarities among *Apibacter* group.

Dataset S4. Distribution of amino acid biosynthesis genes distributions.

Dataset S5. List of sublineage specific gene families.

Dataset S6. List of gene families in the LCA of the three groups.

Dataset S7. Distribution of virulence factors across the three groups.

## Author Contributions

X. Zhang and X. Zhou conceived the experimental design and wrote the manuscript with support from W. Zhang. W. Zhang performed the experiment and bioinformatic analysis. X. Zhang and W. Zhang analyzed data. Q. Su and W. Zhang isolated the *Apibacter* strains used in this work. M. Tang conducted the assembly of the *Apibacter* genomes. X. Zhou supervised the findings of this work.

